# MetaWorks: A flexible, scalable bioinformatic pipeline for high-throughput multi-marker biodiversity assessments

**DOI:** 10.1101/2020.07.14.202960

**Authors:** Teresita M. Porter, Mehrdad Hajibabaei

**Author notes:** Corresponding author: T.M. Porter.

## Abstract

Multi-marker metabarcoding is increasingly being used to generate biodiversity information across different domains of life from microbes to fungi to animals such as for molecular ecology and biomonitoring applications in different sectors from academic research to regulatory agencies and industry. Current popular bioinformatic pipelines support microbial and fungal marker analysis, while ad hoc methods are often used to process animal metabarcode markers from the same study. MetaWorks provides a harmonized processing environment, pipeline, and taxonomic assignment approach for demultiplexed Illumina reads for all biota using a wide range of metabarcoding markers such as 16S, ITS, and COI. A Conda environment is provided to quickly gather most of the programs and dependencies for the pipeline. Several workflows are provided such as: taxonomically assigning exact sequence variants, provides an option to generate operational taxonomic units, and facilitates single-read processing. Pipelines are automated using Snakemake to minimize user intervention and facilitate scalability. All pipelines use the RDP classifier to provide taxonomic assignments with confidence measures. We extend the functionality of the RDP classifier for taxonomically assigning 16S (bacteria), ITS (fungi), and 28S (fungi), to also support COI (animals), rbcL (eukaryotes, land plants, diatoms), 12S (fish), 18S (eukaryotes, diatoms) and ITS (fungi, plants). MetaWorks properly handles ITS by trimming flanking conserved rRNA gene regions as well as protein coding genes by providing two options for removing obvious pseudogenes. MetaWorks is available at https://github.com/terrimporter/MetaWorks along with quick-start instructions using test data, detailed workflow descriptions, and a tutorial for new users.

## Background

Marker gene sequencing, metabarcoding, or metasystematics are interrelated techniques that involves extracting DNA from bulk samples such as soil, water, or mixtures of individuals collected from traps, without having to isolate or identify individual specimens, followed by enrichment of a signature DNA region to identify biological community composition using bioinformatics [1–3]. In microbial ecology to animal biodiversity studies, different signature DNA regions are targeted for their ability to identify target taxa. For example, in prokaryotes, the 16S small subunit (SSU) ribosomal RNA (rRNA) region is often used for genus level taxonomic assignments [4,5]. Other popular markers include cytochrome c oxidase (COI) for animals; ribulose bisphosphate large subunit (rbcL) for plants and diatoms; the internal transcribed spacer (ITS) for fungi and plants; 18S SSU for eukaryotes, arbuscular mycorrhizal fungi, and diatoms; and 12S mitochondrial SSU for fish [6–14].

Existing pipelines such as QIIME2 and DADA2 were initially developed to support the microbial ecology community [15,16]. In comprehensive, multi-trophic, multi-marker studies, there is a need for a pipeline that can handle rRNA genes, spacer regions, as well as protein-coding markers in a single harmonized environment [17,18]. For the ITS region, we needed a pipeline that could remove the conserved flanking rRNA genes as this has been shown to improve taxonomic assignment accuracy [19]. For protein-coding regions, we needed a pipeline that could remove putative pseudogenes [20–23]. We also wanted the ability to generate high quality exact sequence variants (ESVs) for any marker (not just 16S or ITS) for the additional level of genetic and taxonomic resolution SVs can provide [24–26]. For taxonomic assignment, we wanted to use a classifier that would provide a measure of confidence for assignments to reduce false-positive assignments [27–29].

As multi-marker studies are carried out on phylogenetically divergent taxa, such as in biodiversity or trophic studies, there is a need for more generic pipelines where different markers can be analyzed using similar dataflows with 3rd party programs instead of being limited to database-specific pipelines and tools [17,30]. We developed MetaWorks with the following objectives: 1) reproducibility with respect to the computational environment used as well as the pipeline itself, 2) scalability to leverage high performance computer clusters to speed up the analysis of large datasets, 3) naive Bayes classifier support for popular metabarcode markers; and 4) to support marker-specific processing steps such as ITS extraction and pseudogene-removal for protein-coding markers. MetaWorks was designed for data analysts who are comfortable using Linux command-line tools but would like a single harmonized environment and pipeline to process multi-marker metabarcode datasets.

## Implementation & Workflow

### Implementation

MetaWorks is a multi-marker ‘meta’-barcode pipeline that does ‘the works’ by supporting the bioinformatic processing of popular markers including rRNA genes, spacers, and protein coding genes generating taxonomically assigned sequence variants or operational taxonomic units (OTUs). To facilitate reproducibility, scalability, and shareability of workflows we use the Conda package manager to facilitate the download of most programs and dependencies and the Snakemake workflow manager to automate pipelines and utilize computational resources efficiently [31–33]. Snakemake supports re-entrancy and automatic deployment of multiple parallel jobs, both ideal for high performance computing environments where many cores are available to speed up the analysis of large datasets.

We provided instructions on how to install and use Conda in the online documentation. One additional program not available as a Conda package, ORFfinder, may need to be downloaded separately if pseudogene-filtering will be conducted and instructions are provided in the online documentation. MetaWorks is available from https://github.com/terrimporter/MetaWorks and a suite of trained classifiers for taxonomic assignment are also available from GitHub (Table 1). Depending on the DNA metabarcode marker(s) the user will be processing, these can be individually downloaded from GitHub and instructions are provided in the online documentation.

**Table 1.**
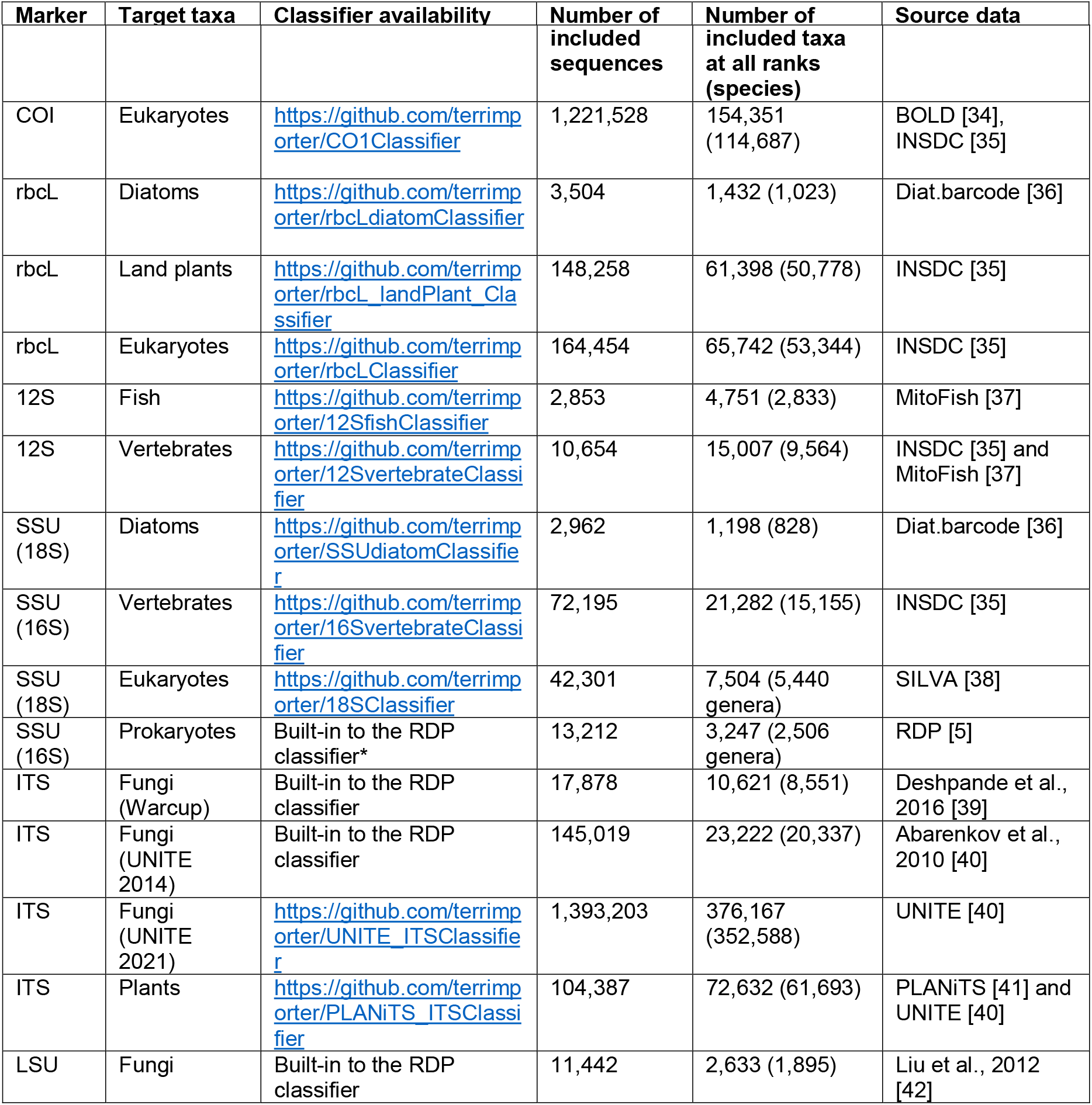
RDP-trained reference sets that can be used with MetaWorks.

### Workflow

The pipeline begins with demultiplexed Illumina paired-end reads as this is the format most often provided by sequencing centres to their clients. Several workflows are available as Snakemake pipelines such as taxonomic assignment of sequence variants (Fig 1), clustering of sequence variants into OTUs, or for processing single reads. For each of these workflows described below, parameter settings for each bioinformatic step can be customized in the config.yaml file. The user also needs to provide a file of primer sequences so we provide a template for the adapters.fasta file as well as a small set of raw Illumina sequences for the COI amplicon that can be used to test the installation. The online documentation provides a ‘quick-start’ example using the provided COI test data. We also provide a more detailed tutorial in the online documentation that walks users through the steps necessary to set up their environment to run the pipeline for the first time, assuming the user has never worked with Conda or Snakemake before.

**Figure 1.**
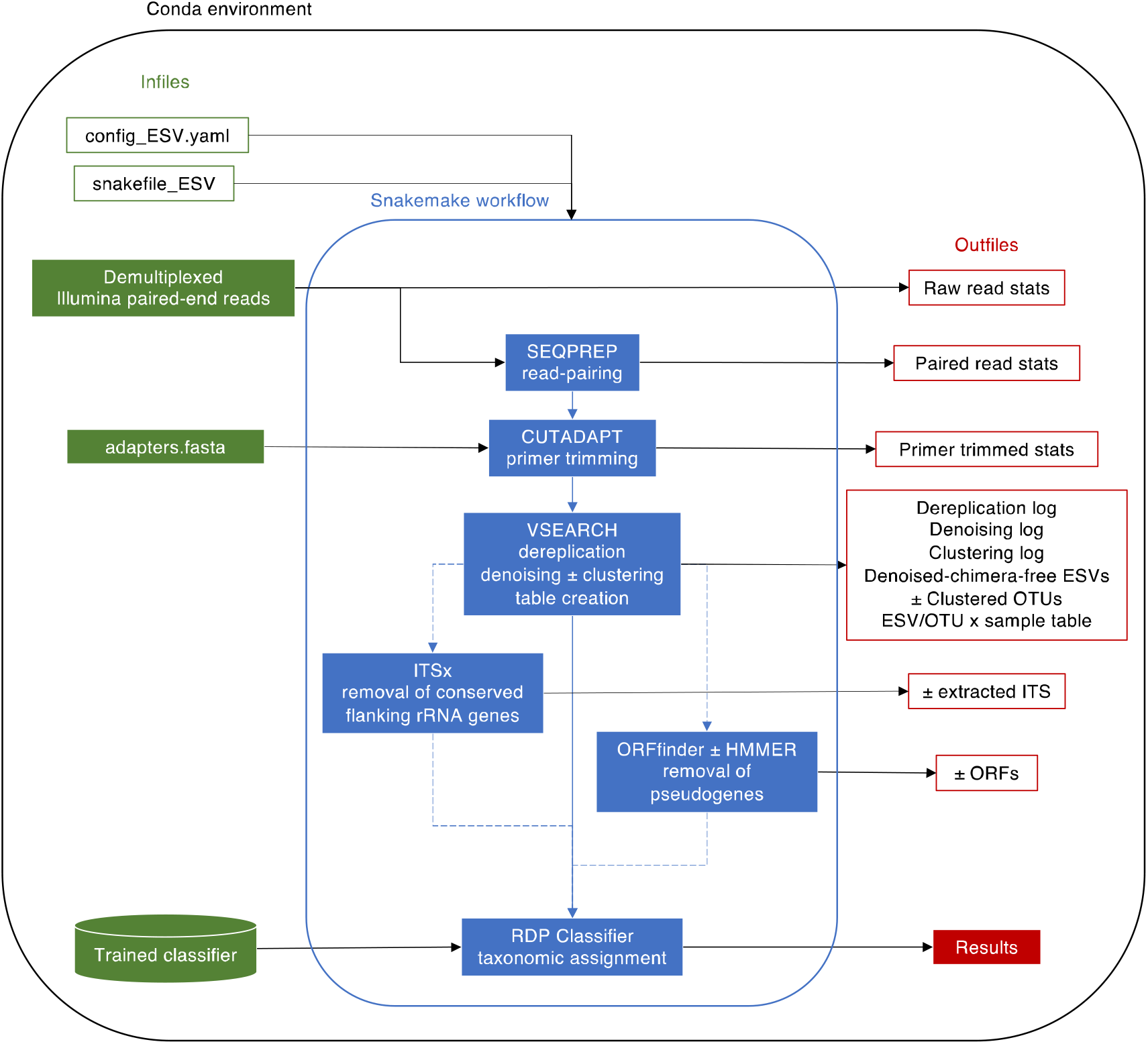
MetaWorks workflow to produce taxonomically assigned sequence variants. To aid reproducibility, a Conda environment is provided (environment.yml). Although multiple Snakemake workflows are provided in MetaWorks, here we show the workflow to produce taxonomically assigned sequence variants. User input is shown in green on the left, the ESV workflow is shown in the middle in blue, and outfiles are shown in red on the right. The results are provided in a comma-separated value (CSV) file and shows each ESV/OTU per sample with read counts and taxonomic assignments. Abbreviations: internal transcribed spacer (ITS) region, open reading frame FASTA-formatted sequences (ORFs).

### Generation and taxonomic assignment of sequence variants

The ESV workflow will run the pipeline shown in Fig 1. In the config_ESV.yaml file, users indicate the path to the directory that contains the demultiplexed Illumina paired-end reads, specify the unique part of filenames to distinguish between samples and reads, and specify the name of the directory that will contain the outfiles. Default settings for each program are provided in the config_ESV.yaml file but these can be customized by the user. SEQPREP was initially chosen for pairing the forward and reverse reads, because the program comes with the option to output the alignments for visual inspection, an option that most read-pairing programs do not have [43]. For SEQPREP read-pairing, users can specify a Phred score quality cutoff, the minimum overlap between the forward and reverse reads, the maximum fraction of mismatches allowed in the overlap region, and the minimum fraction of matching bases in the overlap region. CUTADAPT was chosen for primer-trimming because it is fast and already widely used in the metabarcoding community for this purpose, so most users will likely already be familiar with how this program works [44]. For CUTADAPT, users need to provide a FASTA-formatted primer sequence file (adapters.fasta), they can also specify the minimum sequence length to retain after primer-trimming, a Phred quality score cutoff, the maximum error rate, minimum adapter overlap, and maximum number of ambiguous bases allowed. VSEARCH was chosen to dereplicate reads (retain unique reads) and remove artefactual sequences using the UNOISE3 and UCHIME3 algorithms [45,46]. We chose the open-source VSEARCH program over alternatives because the program can utilize all the available memory on a system, facilitating the analysis of large datasets on high performance computer systems. We prefer the UNOISE3 method for denoising because it performs up to 1,200 faster and uses less memory than other denoising programs [47]. To map read counts to the newly generated denoised-chimera ESVs to create an ESV x sample table, we use the ‘search_exact’ method because it is faster and optimized to find exact matches compared with the ‘usearch_global’ command with the ‘id 1.0’ parameter, but this is just an intermediate step and further filtering of this table is performed by MetaWorks.

If the internal transcribed spacer (ITS) region is analyzed, then the pipeline uses the ITSx program to trim away the flanking conserved rRNA gene sequences so that taxonomic assignment is based solely on the variable spacer region sequences (ITS1 or ITS2) [19]. This step has been shown to improve sensitivity of clustering and taxonomic assignments [19].

If a protein coding marker is being processed, the user can select a pseudogene-removal method in the config_ESV.yaml file. We have previously described two methods for removing putative pseudogenes from DNA barcode and metabarcoding datasets [21]. The NCBI ORFfinder program is used to translate reads into all possible open reading frames (ORFs). The first pseudogene removal method retains the longest ORF for each read, calculates a distribution of ORF lengths, and removes reads with outlier lengths as putative ORFs. The second pseudogene removal method can be used if a hidden Markov model is available and is provided for processing COI arthropods. The longest ORFs are compared to the profile using HMMER available from http://hmmer.org.MetaWorks calculates a distribution of bit scores and removes reads with short outlier bit scores as putative ORFs. Removing noise caused by the sequencing of pseudogenes in metabarcode datasets can help users avoid overe-stimating richness in subsequent analyses, yet this step is not included in the most popular metabarcode pipelines as they were developed to support the analysis of rRNA genes where this is not a problem.

One of the features of MetaWorks, is the use of a single taxonomic assignment method for any metabarcode marker that provides a measure of confidence for taxonomic assignments. We chose the RDP Classifier for this task as this method has a long-history of use in the microbial ecology literature, additionally the classifier can be customized and validated for any metabarcode marker [5]. The RDP classifier calculates k-mer frequencies and uses a naive Bayes method to taxonomically assign unknown query sequences. Bootstrapping is used to provide a measure of statistical support, or repeatability, for each assignment at each rank. We have previously described how this method works compared to the top BLAST hit method [28]. In that comparison, we showed how the RDP classifier is faster than the top BLAST hit method and helps to reduce the rate of false-positive assignments. In studies where erroneously identifying a metabarcode sequence as a potential invasive species or pathogen could lead to alarm, reducing the false-positive assignment rate is critical. We provide a suite of trained classifiers, ready for use with MetaWorks (Table 1). Additionally, we provide the training files so that users can check that key target taxa are present in the reference database, and users are free to use the FASTA-formatted sequence files to create custom BLAST databases for similarity-based searches for data exploration or to build reference sets for subsequent phylogenetic analysis. The final file is a comma-separated value file (results.csv) where the taxonomic assignment for each sequence variant is provided for each sample along with read counts. If an rRNA marker was processed, then the sequence variant sequence is provided in this file; and if a protein coding region was processed, then longest ORF is provided.

### Operational taxonomic units

This pipeline supports the analysis of sequence variants for the additional genetic and taxonomic resolution provided by this level of analysis [24]. Though this method of analysis was initially used to process 16S rRNA genes, studies using ITS and COI have also shown that the analysis of SVs improves the detection of genetic diversity and richness, when assessing beta diversity, both SVs and OTUs tend to recover similar gradients in multivariate analyses [25,26]. Although it has been shown that for many clustering methods sequence order matters and OTU composition can change form one analysis to the next making reproducibility an issue, there are a number of reasons why a user would want to analyze OTUs. For example, it may be more advantageous to work with OTUs instead of sequence variants for network analysis to detect more co-occurrences, for legacy reasons to compare results to previous studies that used OTUs, or to approximate ‘species’ units [48].

After processing raw reads using the snakefile_ESV workflow described in the previous section, users cluster sequence variants into OTUs, a method that combines the benefits of denoising with clustering using a 97% sequence similarity cutoff using the snakefile_OTU workflow [26,49]. This method uses VSEARCH ‘cluster_smallmem’ method to cluster SVs using a 97% sequence similarity cutoff. Settings can be adjusted in the in the config_OTU.yaml file such as pointing to the directory that contains the sequence variants and choosing a classifier for the OTUs.

## Results and Discussion

MetaWorks has already been used in several publications for the Canadian STREAM biomonitoring program, the Government of Canada, Genomics Research and Development Initiative, Metagenomics-based ecosystem biomonitoring (Ecobiomics) project, and by Natural Resources Canada [18,50,51]. The benefits of using an automated, scalable, versioned pipeline for biomonitoring are many-fold, from the ability to share reproducible workflows with collaborators to facilitate the re-analysis of data as more samples are collected from year-to-year. Below, we describe three MetaWorks use-cases in more detail below.

### Use case 1

As a part of the Canadian STREAM biomonitoring Initiative, the MetaWorks pipeline has been used to process macroinvertebrates COI metabarcodes surveyed from stream sites across Canada [50]. One feature of this project is the quick 1-2 month turn-around time from sampling through to the production of watershed biodiversity reports. This is an improvement over reports generated using conventional morphology-based methods that would normally take 6 - 12 months to produce. The use of a consistent bioinformatics workflow to process metabarcodes has played a key role in the reproducibility, scalability, and throughput to facilitate timely reporting [52]. Generally, samples are processed in batches of 96 per sequencing run then later split into custom reports for stakeholders, processing about 500 samples per year. One feature of these reports are the taxonomic assignments made using the naive Bayesian classifier that can be filtered for identifications to the species, genus, and family ranks with 99% accuracy assuming the query is represented in the underlying sequence database [28]. This is in contrast with the use of more traditional methods for taxonomic assignment, where taxa are routinely missed during subsampling and taxa detected by primary analysts and auditors may differ by up to 30% [53]. This use-case shows how MetaWorks can be used to create taxon lists for large-scale biodiversity monitoring of streams across Canada.

### Use case 2

Also as a part of the STREAM project described above, MetaWorks results were to analyze sequence variants from diatoms (rbcL) and arthropods (COI) sampled within and across sites of varying water quality [54]. Using the MetaWorks pipeline, two different protein-coding markers were bioinformatically processed in two runs. The first run processed the rbcL marker, using a mixture of 5 different primers in a single adapters.fasta file, and pseudogenes were removed from this dataset using the simple ORFfinder method [21]. The second run processed three COI amplicons, each targeting an approximately 200 bp length of the COI barcoding region using 6 different primers in a single adapters.fasta file, and pseudogenes were removed from this dataset using the ORFfinder+HMMER method since a COI arthropod HMM model was available [21]. The study reported a diversity assessment across sites of varying water quality using richness, effective richness, and beta diversity. Additionally, the taxonomic assignments generated from MetaWorks were used to obtain resource-consumer relationships from a global database of biotic interactions (GloBI) so that community stability using trophic and network measures could be assessed across sites with varying water quality [55]. This use-case shows how MetaWorks can handle a variety of protein-coding markers for trophic and network analyses to facilitate ecological assessments of freshwater condition.

### Use case 3

As a part of collaborative work with Environment and Climate Change Canada, MetaWorks was used to assess macroinvertebrate and (non-macroinvertebrate) eukaryote taxa in an urban harbour using COI and 18S rRNA [56]. Using the MetaWorks pipeline, COI metabarcodes were identified down to species rank with 99% accuracy and 18S metabarcodes were identified to genus rank with 80% accuracy using a custom-trained classifier based on the SILVA 18S release 138 [38]. In this study, conventional macroinvertebrate sampling for assessing water quality in Toronto Harbour was compared with metabarcoding methods. COI metabarcoding was found to detect more diversity at a finer level of taxonomic resolution compared with conventional approaches and was able to distinguish sites with particularly high levels of sediment contaminants. Additionally, the use of a multi-marker approach allowed microscopic eukaryote diversity to be sampled at the same time from the same samples, producing indicators that responded to gradients in both sediment contaminants and water physical-chemical features. This use-case illustrates how MetaWorks can facilitate the application of multi-marker metabarcoding approaches that target different domains of life.

As demonstrated in the above examples, MetaWorks supports a wide range of analysis scenarios from metabarcoding data. We envision that MetaWorks will aid broader user communities and fill a need in multi-marker metabarcoding studies that target taxa from multiple different domains of life, to provide a unified processing environment, pipeline, and taxonomic assignment approach for each marker from ribosomal RNA genes, spacers, or protein coding genes. QIIME2 is perhaps the most popular and comprehensive platform for such work, but to date, focuses on processing mainly prokaryote and fungal datasets [16]. As of yet, MetaWorks is the only bioinformatic pipeline that can handle rRNA genes but that also integrates special processing steps to handle ITS spacers as well as filter out obvious pseudogenes in protein coding markers such as COI.

There has been a lot of activity with respect to building new bioinformatic tools to handle COI metabarcodes. Recent work, such as the BOLDigger program, has attempted to make the BOLD identification engine more suitable for identifying large batches of COI metabaracodes and has both GUI and command-line interfaces for efficient sample processing [57]. A new program, called NUMTdumper, has been developed as a stand-alone program meant to be incorporated into bioinformatic pipelines [20]. NUMTdumper provides a method to screen for NuMTs based on read counts while acknowledging the trade-offs between removing all possible NuMTs while erroneously removing genuine reads. An R package called ‘coil’ has also recently been developed that will place COI barcode and metabarcode sequences in frame using profile HMM analysis [58]. MetaWorks aims to extend the COI metabarcode toolkit that provides a harmonized environment where data from other organismal markers in multi-marker, multi-trophic studies can also be analyzed.

## Conclusion

MetaWorks is provided as free and open software that is versioned, can be easily deployed in a Conda environment, and is supported by a suite of classifiers for popular metabarcoding markers. The software comes with a small set of raw data and a quick-start pipeline to help users gain experience quickly. There is extensive online documentation available at the link above included detailed explanations of the pipeline, available workflows, and a tutorial for new users who have never used Conda or Snakemake before. MetaWorks generates a CSV file that lists all sequence clusters, for each sample, with associated read counts, taxonomic assignments, and bootstrap support values. Numerous statistics and log files are also provided so that users can track the number of reads that pass each major bioinformatic step. Given the current use of MetaWorks by large-scale national initiatives such as STREAM and Ecobiomics, we foresee additional developments and enhancements. Future improvements include the development of additional HMM models for pseudogene filtering, updated and additional classifiers for taxonomic assignment, and support for processing larger jobs both on HPCs and in a cloud environment. We welcome suggestions and potential collaborative work to further advance this pipeline for the scientific community.

## Availability and requirements

MetaWorks runs at the command-line in a Conda environment on a Linux operating system. Miniconda can be downloaded from https://repo.continuum.io/miniconda/Miniconda3-latest-Linux-x86_64.sh. If pseudogene filtering will be performed ORFfinder can be downloaded from the NCBI ftp site at ftp://ftp.ncbi.nlm.nih.gov/genomes/TOOLS/ORFfinder/linux-i64/ORFfinder.gz. MetaWorks workflows, quick-start example, and a tutorial for new users is available from GitHub at https://github.com/terrimporter/MetaWorks. Additional trained classifiers that can be used by MetaWorks for taxonomic assignment are available from GitHub at https://github.com/terrimporter.

## Funding

MH received funding from Genome Canada and Ontario Genomics for the Sequencing the Rivers for Environmental Assessment and Monitoring (STREAM) project. TMP received funding from the Government of Canada through the Genomics Research and Development Initiative (GRDI), Metagenomics-based ecosystem biomonitoring (Ecobiomics) project.

## Authors’ contributions

TP and MH conceived of the work. TP designed and wrote the pipeline. All the authors helped to write the manuscript, approve the submitted version, and agree to be personally accountable for the author’s own contributions and to ensure that questions related to the accuracy or integrity of any part of the work, even ones in which the author was not personally involved, are appropriately investigated, resolved, and the resolution documented in the literature.

## Acknowledgements

We would like to thank Josip Rudar, Katie M. McGee, Chloe V. Robinson, Victoria C. Maitland, and Michael T.G. Wright from the Hajibabaei lab for helpful discussions and testing the pipeline with various datasets.

## Notes

https://github.com/terrimporter/MetaWorks

### Competing Interest Statement

The authors have declared no competing interest.

### Summary of Updates

Updated to reference newly created classifiers and workflows since the last version.

